# Enhancing experimentally the structural heterogeneity of forests increase soil fungal diversity but functional lifestyles in contrasting ways

**DOI:** 10.1101/2025.08.29.673114

**Authors:** Claus Bässler, Petr Baldrian, Jörg Müller, Franz-Sebastian Krah, Mirjana Bevanda, Pia Bradler, Antonio Castaneda-Gomez, Benjamin M. Delory, Sebastian Dittrich, Nico Eisenhauer, Andreas Fichtner, Akira S. Mori, Minagi Naka, Goddert von Oheimb, Luisa Pflumm, Martin Wegmann, Vendula Brabcová

## Abstract

Fungal communities in soils are highly diverse both in species and functions forming a major backbone of forest ecosystems. Recent observational high-throughput-sequencing studies have shown that fungal diversity is correlated with resource availability and climate across different spatial scales. However, the underlying mechanisms remain poorly understood. Across Germany, we experimentally manipulated 11 typically homogeneous, broadleaf production forests to increase their between-patch-heterogeneity (ESBC) and compared them with a control forest. In specific, we enhanced light availability via canopy openness and deadwood resources in the ESBC treatments. Fungal communities were determined by metabarcoding at 234 patches and analysed using a novel meta-analytical approach for pairwise comparisons of taxonomic and phylogenetic diversity along Hill numbers. We hypothesized that γ-diversity is primarily driven by β-diversity increasing with canopy mediated microclimate variability and secondarily by α-diversity increasing with resource availability. Furthermore, we expected an increase in γ-diversity by unique phylogenetic lineages supporting the *insurance hypothesis*. Our results showed a significant increase in γ-diversity in ESBC forests, first by α- and second by β-diversity, both of which were influenced mainly by microclimate and not resource availability. The increase of phylogenetic diversity with ESBC was weak indicating functional similarity of species. Analysis of symbiotic, saprotrophic and parasitic fungal assemblages revealed contrasting effects of resource availability and microclimate across the scales. As in the UN Decade of Ecosystem Restoration many forest managers aim to increase the heterogeneity of forests and are simultaneously face rising tree mortality, our study provides first robust empirical evidence for the varying effects of forest gaps and deadwood on fungal diversity across α-, β-, γ-scales for this major kingdom.

## Introduction

Fungi form a hyperdiverse kingdom and have evolved a wide range of functional lifestyles, playing key roles in many ecosystem processes (Baldrian 2017). Symbiotic fungi contribute substantially to the productivity of ecosystems via the provisioning of water and nutrients to host species (van der Heijden et al. 2006) and mediating the flow of photosynthetic carbon into soil (Hawkins et al. 2023). Saprotrophic fungi belong to the most efficient organisms to break down organic matter (Floudas et al. 2012), while fungal parasites exploit living resources in manifold ways, thereby affecting their diversity and structure (Runnel et al. 2025).

In contrast to plants and animals, the hidden living of most fungi species prevented long time the exploration of their diversity patterns. During the last decades, high throughput sequencing boosted efforts to study fungal diversity patterns, particularly at very small and very large scale (Baldrian 2017, Nilsson et al. 2019). Today we know that soil fungal diversity shows a biogeographic patterning, mostly shaped by macroclimate (Větrovský et al. 2019, Mikryukov et al. 2023, Abrego et al. 2024). Numerous studies on α- and β-diversity at smaller spatial scales revealed that particularly resource related variables (e.g., nutrients vegetation structure) explain soil fungal diversity (e.g., Glassman et al. 2017, Yang et al. 2017, Odriozola et al. 2021). However, from the existing studies, we identify three important knowledge gaps, which prevent further mechanistic insights: First, most studies are observational, which hampers disentangling resource related from abiotic factors, particularly at larger spatial scale. Second, studies are conducted either on large macroecological scales or local α-scales. Therefore, despite the overwhelming importance of fungi in many ecosystems, e.g., forests, replicated studies across scale relevant for landscape management are lacking. Hence, the relative importance of α- and β-diversity and related drivers are still unknown. Third, the link between diversity and ecosystem stability has been subject to numerous studies the last decades (Tilman et al. 2006). The basic idea is that ecosystem stability is related to a broad range of species that respond differently with environmental fluctuations (e.g., *insurance hypothesis*, Yachi and Loreau 1999 or, portfolio effect, Lhomme and Winkel 2002). Indeed, numbers of studies demonstrated that a local diversity loss (α-scale) is related to a loss of ecosystem functioning and thus, reduce stability (Cardinale et al. 2012, Naeem et al. 2012), confirmed also for soil fungi (Liu et al. 2022). However, a recent global meta-analysis has shown that the impact of various anthropogenic stressors on different scales of diversity can vary considerable even for microbes (Keck et al. 2025). Therefore, current evidence of the relationship between α-diversity and ecosystem stability from local studies cannot simply be translated to larger spatial scale (Wang and Loreau 2016). With this study, we, thus, aim to investigate overall soil fungal diversity patterns and those of important functional groups at the different scales.

We enhanced experimentally the between-patch heterogeneity (ESBC) of 11 forests by creating gaps and deadwood throughout Germany and selected for each of them a comparable homogenous control forest (Müller et al. 2023) (Fig. 1). We identified soil fungal communities at each patch via high-throughput sequencing and assigned the OTUs (termed ‘species’ in the following) to the three major functional groups; symbionts, saprotrophs and, parasites. We applied a newly developed meta-analytic approach, which allows considering undetected species, the entangling of α-, β- and γ-diversity along the Hill numbers and, simultaneously taxonomic and phylogenetic diversity in a unified approach.

**Fig. 1.**
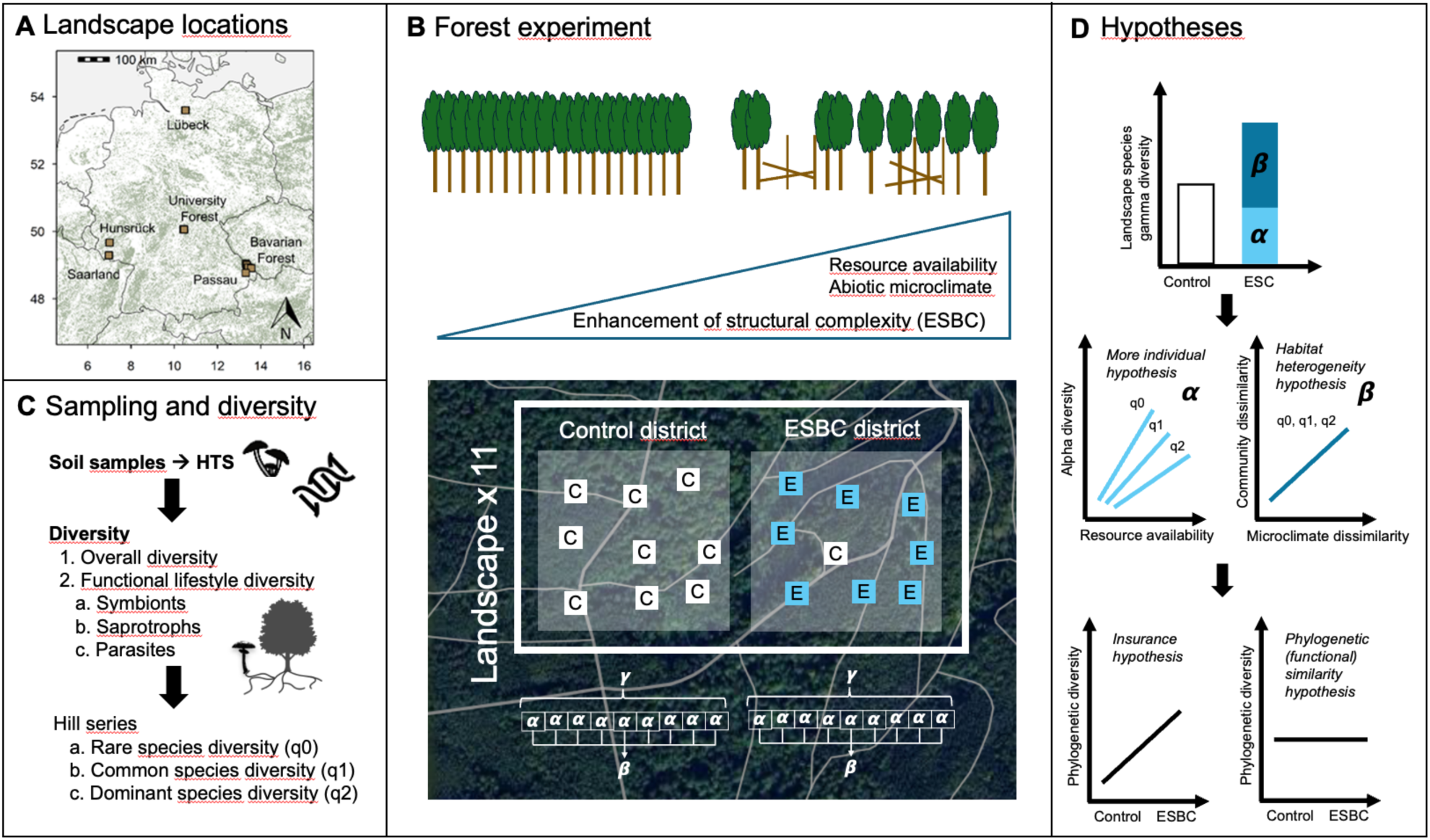
**A.** Location of the 11 experimental landscapes within Germany. **B.** Experimental setting: in each landscape, we set up a control and a treatment (enhancement of structural complexity = ESBC) district. ESBC treatment is characterized by two main structural axes (deadwood (resource) enrichment and canopy mediated abiotic microclimate). **C.** Sampling of soil fungal communities and diversity measures used. **D.** Hypotheses derived from theory and empirical studies (see Introduction).

We hypothesized that landscape γ-diversity is higher in ESBC than in control forests (Fig. 1). We further hypothesized that this effect is mainly caused by an increase in β-diversity as across studies and scales, consistent effects of the environment on soil fungal communities has been shown, while α-effects were partly weak or inconsistent (e.g., Baldrian 2017, Odriozola et al. 2024). Opening the canopy change suddenly and strongly environmental conditions (i.e., microclimate) in forest ecosystems (Thom et al. 2020) and thus, increase abiotic heterogeneity. According to the *habitat heterogeneity hypothesis* (MacArthur and MacArthur 1961), we, thus, expect an increase of β-diversity fueling γ-diversity due to canopy cover differences among the control versus ESBC patches (Štursová et al. 2014). However, ESBC patches are enriched with deadwood, and it has been shown that availability of deadwood increases soil fertility and improve soil physical properties (Sokołowski et al. 2025). Thus, increased chemical energy might contribute to explain γ-diversity via an increase in fungal α-diversity which would be in line with the *more individual hypothesis* (Sterkenburg et al. 2015, Storch et al. 2018). Here, effects might be stronger for q0 (rare species diversity) as increased resource availability might allow the coexistence of rare species (Thomas 1994) (Fig. 1). Finally, an increase of unique phylogenetic lineages with ESBC would support the *insurance hypothesis* (Yachi & Loreau 1999) under the assumption that different phylogenetic lineages contribute to ecosystem function in complementary ways (Srivastava et al. 2012, but see, Cadotte et al. 2017). In contrast, under this assumption, a lack or weak response of phylogenetic diversity would support the functional similarity hypothesis (Eisenhauer et al. 2023). We hypothesize that (i) overall soil fungal γ-diversity increases with ESBC and this effect is mainly caused by an increase of β-diversity due to changes in canopy cover; (ii) α-diversity contributes to explain an increase of γ-diversity with ESBC and this effect is mainly attributable to the enrichment of deadwood. To further illuminate the functional consequences of ESBC (i.e. inferences about ecosystem stability), we consider phylogenetic diversity and evaluated the response of the main fungal functional lifestyles (symbionts, saprotrophs and parasites).

## Methods

### Study area and study design

Within each of the 11 sites (landscapes), the ESBC and control district each consists of nine different patches with ca. 10-20 ha extend. In three sites, we considered three patches in addition (15 patches). At each site, one district was subjected to silvicultural interventions to manipulate the variation in canopy cover and deadwood amount across patches (termed ESBC treatment). We manipulated ca. 30% of trees within the 50 by 50 m patches and left snags, logs, stumps or combinations in either aggregated (canopy open) or distributed (still closed canopy) spatial arrangement (Fig. 1). In the three sites with the 15 patches, we considered in addition total tree removal, left crowns and created habitat trees also in aggregated and distributed spatial arrangement (see Kacic et al. 2024 for details on the structural effects of ESBC). The ESBC manipulation took place from 2016-2019 depending on the site (Müller et al. 2023). Overall, the ESBC treatment reflects within landscape structural heterogeneity. The second pair of each site with 9 respectively 15 patches acted as control. These are characterized by uniform thinning operations approximately every 10 years, as is typical for production forests of this age (Uhl et al. 2025). Also here, a comparable proportion of trees (∼30%) as in the ESBC treatment is removed in the form of thinning. However, no deadwood from harvest residues remains on the site and no sustained openings in the canopy are produced. This design offers realistic comparisons between current production and ESBC forests with replications of landscapes covering variability in climate conditions of temperate broadleaf forests in Europe.

### Sampling soil fungal communities

Soil sampling followed a standardized protocol across all plots using a plastic soil corer (4.5 cm diameter). The reference point in each patch was the approximate central geometric point. A central core was collected 1 m north of the reference point, followed by four additional cores taken 2 m away in the cardinal directions (N, E, S, W), forming a cross-shaped layout (Martinović et al. 2021). Surface litter (undecomposed leaves or needles) was removed prior to coring. Each core was extracted to a depth of approximately 15 cm. The five cores from each patch were combined into a composite sample, with only the upper 10 cm of soil retained. The material was thoroughly mixed, homogenized, sieved through a 5 mm mesh, and manually cleared of visible roots and remaining litter. An aliquot of the homogenized soil was immediately frozen, freeze-dried, and stored at −20 °C until further analysis. Samples were kept at 4 °C prior to processing and processed within 48 hours of collection. At each site, photographs of the forest floor, characteristic vegetation, and canopy were taken for documentation.

Total genomic DNA was extracted in triplicate from 150 mg of freeze-dried soil per sample using a modified Miller method (Sagova-Mareckova et al. 2008). This method involves phenol/chloroform extraction with bead-beating using a FastPrep-24 instrument (MP Biomedicals, Santa Ana, USA) at 5 m s⁻¹ for 2 × 30 seconds. DNA extraction was followed by purification using the GeneClean Turbo Kit (MP Biomedicals). The pooled extracted DNA was quality-checked and quantified with a NanoDrop ND-1000 spectrophotometer (Thermo Fisher Scientific) stored at −20 °C for further analyses.

The fungal community structure was addressed using barcoded gITS7 and ITS4 to amplify fungal ITS2 region (Baldrian et al. 2016). To minimize PCR bias, polymerase chain reaction (PCR) was performed in triplicate for each DNA sample. Each PCR contained 5 µl of 5× Q5 reaction buffer, 1.5 µl of BSA (10 mg ml-1), 1 µl of each primer (0.01 mM), 0.5 µl of PCR Nucleotide Mix (10 mM each), 0.25 µl of Q5 High Fidelity DNA polymerase (2 U µl -1, New England Biolabs, Inc.), 5 µl of 5× Q5 HighGC Enhancer and 1 µl of template DNA (approx. 50 ng µl-1). Cycling conditions were 98 °C for 30 sec, followed by 30 cycles of 94 °C for 10 sec, 56 °C for 30 sec, and 72 °C for 30 sec, and a final extension at 72 °C for 2 min. Negative and positive amplification controls were used to avoid contamination errors and to prove the efficiency of the PCR, but these controls were not sequenced within this project. PCR triplicate reaction products were pooled and purified (MinElute PCR Purification Kit, Quiagen). Amplicon libraries were prepared with the TruSeq DNA PCR-Free Kit LP (Illumina) and sequenced in house on the Illumina MiSeq (2 × 250-base reads).

Amplicon sequencing data were processed using the pipeline SEED 2.1.3 (Vetrovsky and Baldrian 2013). Briefly, paired-end reads were merged using fastq-join (Aronesty 2013). The ITS2 region was extracted using ITS Extractor 1.0.11 (Bengtsson-Palme et al. 2013) before processing. Chimeric sequences were identified and removed using Usearch 11.0.667 (Edgar 2010), and remaining sequences were clustered into Operational Taxonomic Units (OTUs) using the UPARSE algorithm (Edgar 2013) at a 97% similarity threshold. The most abundant sequence in each OTU cluster was used as the representative for taxonomic assignment using BLASTn against the UNITE 10 database (Abarenkov et al. 2024). Non-fungal sequences were excluded. Fungal OTUs were further classified into functional lifestyles using FungalTraits (Põlme et al. 2020). Sequencing data have been deposited in the NCBI SRA database under BioProject accession number PRJNA1232019.

### Phylogenetic tree

Statistical analyses were conducted in R (version 4.5.0) (R Core Team 2025).

We build phylogenetic hypotheses for fungi. We used a backbone guide tree resolved on the genus level and added missing genera from the dataset in the guide tree based on higher taxonomic levels. We then used the ITS consensus sequences (fasta files) to infer within-genera phylogenetic trees. We replaced the genus tip labels with the within-genera phylogenetic trees. As guide trees for fungi, we used the backbone guide tree by (Tedersoo et al. 2018). Following a published approach (Kortmann et al. 2025), the within-genera phylogenies were based on a multiple sequence alignment (MSA) using the AlignSeq function from the DECIPHER-package. The AlignSeq method performed better than most alignment methods in simulations and real data (Wright 2015). Based on the MSAs, we built the phylogenetic trees using the Neighbor-Joining method implemented in TreeLine. We chose this method, balancing speed, accuracy, and relatively short evolutionary time scales within genera. The resulting trees still contained polytomies, which we randomly resolved 100 times.

We then used the Grafen method to compute branch lengths (Grafen 1989) due to the enormous size of the tree. We tested the variability among the 100 trees (randomly resolved polytonies and subsequent branch length estimation) for our ecological inferences. This sensitivity analysis yielded similar results (data not shown) and we, therefore, present the results for the first tree from the list of the produced 100 (see Appendix S1 for the used phylogenetic tree in newick format). Note that in phylogenetically informed ecological analyses, the topology of the phylogeny affects ecological inference, whereas variations in branch lengths did not have such effects (Cadotte 2015). Note furthermore that, we considered only OTUs in the phylogeny for which we have the information for the taxonomic class (“mycetes”). Thus, from the overall OTU number of the complete data set (19,323 after singeltons correction, see below), we considered finally 12,500 OTUs in our analysis of the overall fungal diversity (taxonomic and phylogenetic α-, β- and γ-diversity). However, sensitivity analysis of the full data set for overall taxonomic diversity yielded similar results (Fig. S1). The functional lifestyles specific analyses were based on the full data set for which, we have lifestyle information as here, we ignore phylogenetic diversity, which is meaningful only for the overall fungal diversity analysis (see below).

### Statistical analysis

#### Singletons correction

We first used the approach of (Chiu and Chao 2016) to estimate the true number of singletons, and we modified the raw community matrix accordingly. Specifically, this approach modifies an OTU table by randomly setting singleton entries to zero such that the number of remaining singletons per sample is equal to the number of singletons estimated by Chiu & Chao’s method for sequencing error estimation. As this procedure involves a resampling approach, multiple runs (i.e., modified community matrices) should ideally be considered in the subsequent analysis. However, testing different resampled community matrices did not alter the inference of our final models. Therefore, for simplicity and reproducibility, we used a single representative community matrix after singleton correction, applying a fixed seed in R.

The final overall community matrix consisted of 19.323 OTUs across 234 patches. We further filtered this community matrix into three community matrices representing the main fungal functional lifestyles. Symbionts were represented by 3,468 OTUs from which >92% were ectomycorrhizal, the remaining mostly other mycorrhizal types (e.g., arbuscular mycorrhiza) and very few animal symbionts (<0.3%). Saprotrophic fungi were represented by 6,577 OTUs, from which ca. 80% were litter, soil and wood saprotrophic fungi, the remaining OTUs were mostly unspecified saprotrophic fungi, and a few belong to other decomposer (e.g., dung saprotrophic). Parasites were represented by 2,034 OTUs from which >98% were animal, plant and fungal parasites, the remaining few belonged to unspecified-, algal- or protistan parasites. Note that not for all OTUs, we have reliable genera information, which explains the difference between the overall number of OTUs and the OTU sum across lifestyles. For simplicity, we term OTUs in the following as ‘species’.

#### Hill numbers

We used our diversity measures as Hill numbers. The Hill numbers can be understood as the effective diversities of communities (Hill 1973). The Hill numbers framework unites established diversity indices that are differently sensitive to rare, common and dominant species. The adjustable parameter q indicates how sensitive the index is to species abundances (Chao et al. 2014). The Hill number of order q = 0 corresponds to the presence/absence of species and thus gives more weight to species in the community with low frequencies such as rare species. This measure reflects species richness. The Hill number of order q = 1 corresponds to Shannon diversity, representing the effective number of common species, while the Hill number of order q = 2 corresponds to inverse Simpson diversity, representing the effective number of dominant species giving weight to the most abundant species. For β-diversity, *q* = 0, 1, and 2 correspond to the Sørensen, the Horn, and the Morisita-Horn indices, respectively.

Hill-Chao numbers, as a generalisation of Hill numbers to phylogenetic diversity, enable a direct comparison of taxonomic and phylogenetic diversity in the same unit of species- or lineage equivalents (Chao et al. 2021): Taxonomic diversity (TD) quantifies the effective number of equally-abundant species. For phylogenetic diversity (PD), we used mean-PD (PD divided by tree depth), which quantifies the effective number of equally-divergent lineages, thus considering species’ evolutionary histories. Hill-Chao numbers are then formulated based on the species abundance distribution using adapted functions from the R packages iNEXT.3D (Chao et al. 2021) and iNEXT.beta3D (Chao et al. 2023)

#### Calculation of α-, β- and γ-diversity diversity across landscapes (meta-analysis)

We calculated the effective number of species and lineages for Hill numbers (q = 0, 1, 2) to allow directly the effects of taxonomic and phylogenetic diversity at the same unit (see above). These diversity measures of soil fungal diversity and corresponding confidence intervals were calculated for each of the 22 districts, using the iNEXT.3D-package and the included bootstrapping approach. γ-diversity (at the landscape scale) was decomposed into α-(patch scale) and multiplicative β-diversity (among patches) using iNEXT.beta3D (version 2.0.8) (Chao et al. 2023). We used the 1-S transformation, a normalized dissimilarity measure equivalent to Jaccard-type turnover, mapping β-diversity to the range of 0 indicating identical species composition among patches to 1 representing completely distinct communities. Standardization also controls for variation in patch numbers of sites (9 and 15, see above). For comparison between ESBC and control districts, we calculated differences in diversity metrics for each of the 11 pairs with positive values indicating higher diversity in the ESBC district. Differences from the 11 forest pairs were combined by meta-analysis propagating site-wise bootstrapping confidence intervals to the aggregated metric. Confidence intervals that exclude the value 0 indicate significant differences. As observed diversity data can be sensitive to sampling effort and sample completeness (Chao and Jost 2012), we standardized diversity estimates to a common coverage level by rarefaction and extrapolation for the overall analysis (SC=0.981) and the functional lifestyle specific analysis (SC=0.988) (Chao et al. 2020). All computation and graphics of the meta-analyses were performed using the new function, iNEXTmeta_beta, adapted from iNEXT.beta3D (Chao et al. 2023). The function is currently available at: https://github.com/AnneChao/iNEXT.meta

#### Linear mixed modelling of α- and β-diversity versus deadwood volume and canopy openness

To inform the response of α- and β-diversity in relation to the ESBC treatments from the meta-analysis, we run linear mixed effects models for overall fungi (taxonomic and phylogenetic diversity) and the different functional lifestyles (taxonomic diversity) using deadwood amount and canopy openness as predictors. Deadwood amount (m^3^*ha^-1^) was measured at each patch by an inventory of all deadwood objects > 7cm in the patch. Canopy openness was measured by LiDAR via Drone flights in the year of fungi sampling using the inverse penetration ratio at 6m above ground (Müller and Brandl 2009). To estimate taxonomic and phylogenetic α-diversity, we used the ‘estmate3D’ function form the iNEXT.3D-package using the same sample coverage levels used for the meta-analysis (see above). Then, we applied negative binomial models (function ‘glmmTMB’ within the glmmTMB-package) (Brooks et al. 2017) using log_10_-transformed deadwood volume and canopy penetration ratio as predictors and site as a random effect. As there might be additional, more subtle environmental variation among patches not adequately reflected by deadwood volume and canopy openness, we additionally used unweighted mean Ellenberg indicator values (Ellenberg 1991) from patch related plant releves. These measures have been shown to be integrative in reflecting subtle environmental conditions (Ewald 2003, Müller and Brandl 2009). We used the values reflecting temperature (L), moisture (F), soil reaction (R, ∼pH-value), nitrogen (N) and continentality (K) but not light (L, as measured directly and used already as fixed effect, see above). To reduce the number of predictors, we subjected the Ellenberg values to a PCA. The first two axes explained ca. 90% of the variability and were represented by the R-(∼pH-value) and nitrogen (N) value respectively. Thus, we used the first two dimension (scores) from the PCA as covariates in our models. Due to low variation within phylogenetic diversity, particularly for q1 and q2 (e.g., for q2; min=1.78, max=3.53, median=2.62), the negative binomial model did not converge. In contrast, using a linear mixed effects model (function ‘lmer’ within the Lme4-package) (Bates et al. 2015) with gaussian distribution produced robust estimates. Note further, phylogenetic diversity for all Hill numbers were normally distributed and thus, the distribution assumption is valid. To compute pairwise β-diversity indices (distance matrices) as a measure of dissimilarity for all study plots, we utilized the function ‘iNEXTbeta3D_pair3D’ adapted from the iNEXT.beta3D-package (see above). Then, we fitted linear mixed effects models (function ‘lmer’, see above) and used the pairwise difference in deadwood volume and canopy openness as fixed factors and site as a random effect to capture both within-site and among-site dependence (Harrison et al. 2018, Chao et al. 2024). As for the α-models, we used the pairwise difference in Ellenberg values calculated from a gower distance as a covariate in the models. Further, in the β-diversity models, we considered the spatial distance among patches based on projected coordinates as predictor. With our spatially explicit design, we entangled environment from space. Thus, considering space in our β-models allows further inferences about assembly mechanisms (e.g., dispersal, drift) explaining diversity differences between control and ESBC treatments (Mori et al. 2018). In all mixed effects models, we estimated the effect sizes for deadwood amount, canopy openness and the covariates separately for the control and the ESBC treatments to be consistent with the approach of the meta-analysis (see above). All models were checked for colinearity among predictors (|r|<0.7) (Dormann et al. 2013) and via variance inflation factors (all <5). We reported z-values as a standardized metrics of effect sizes, significance level (* p<0.05, ** p<0.01, *** p<0.001), the marginal (R^2^m) and the conditional (R^2^c) R-square.

## Results

### Overall soil fungal diversity

Our analysis across the 11 pairs showed a significant increase in taxonomic and phylogenetic γ-diversity of overall fungi in ESBC treatments (Fig. 2A). Effects are considerably stronger for taxonomic diversity and for q0 compared to q1 and q2 (Fig. 2A). Gain of species due to the ESBC treatment at γ-scale was ∼ 95 (83-107, see also Fig. S2-7).

**Fig 2.**
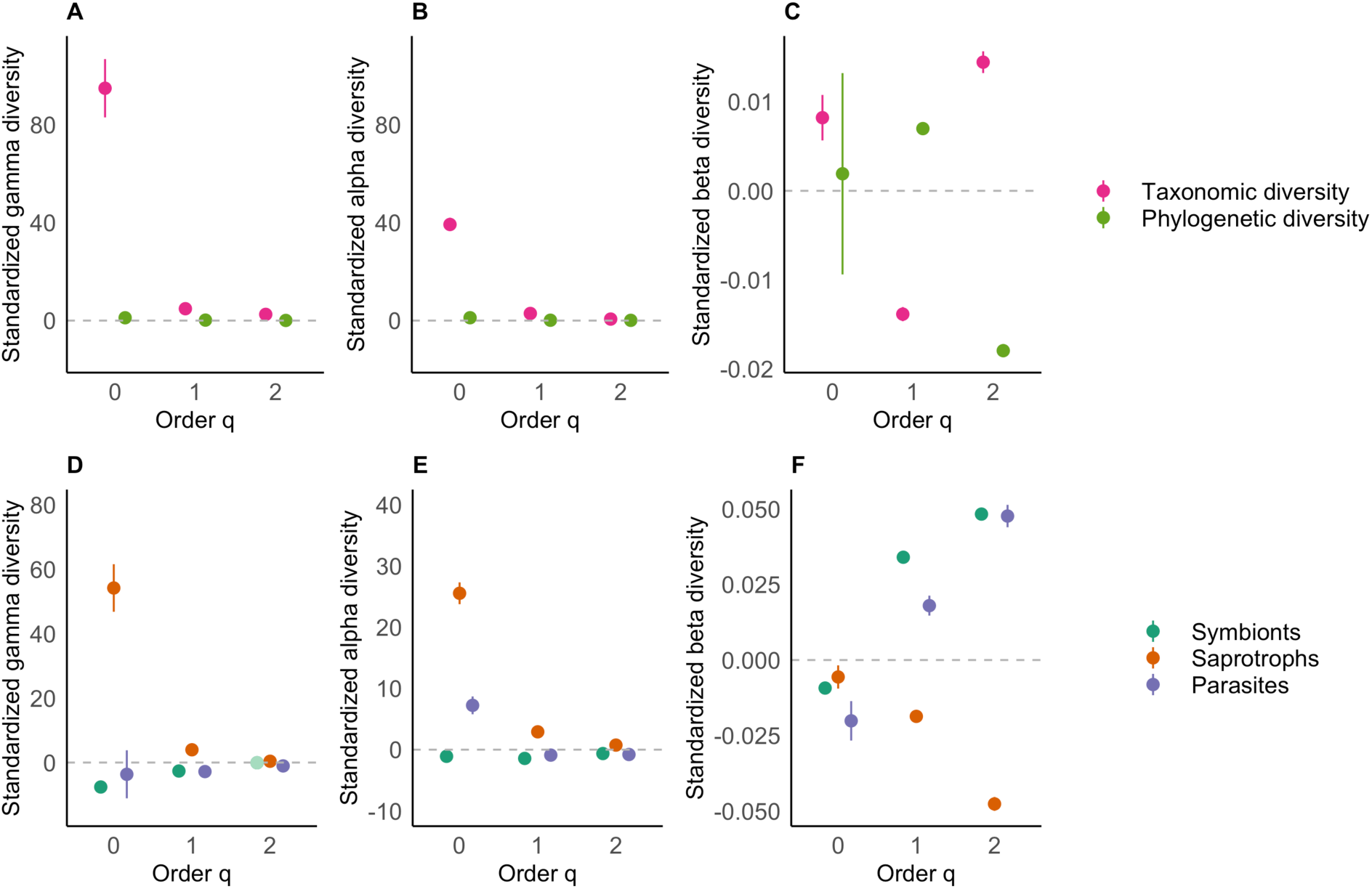
Results from the meta-analysis across 11 experimental landscapes for the overall taxonomic and phylogenetic fungal diversity. (A: γ-diversity, B: α-diversity, C: β-diversity) and separately for the fungal functional lifestyles (D: lifestyle specific γ-diversity, D: lifestyle specific α-diversity, F: lifestyle specific β-diversity). Points indicate the predicted means, and error bars denote the 95% confidence interval. The standardised diversity difference is calculated between each pair of the control and ESBC treatment in directly comparable species-equivalent units for γ- and α-diversity and for the two diversity facets (taxonomic and phylogenetic) and three diversity orders (q = 0, q = 1 and q = 2). A Jaccard-type turnover transformation (1-S) of multiplicative β-diversity was applied to correct for the different numbers of assemblages (i.e. 9 or 15) sampled in the different experimental landscapes. Positive differences (effects) indicate an increase in diversity from control to ESBC districts. Effects are significant when the confidence interval does not include zero.

For α-diversity, we found a similar pattern as for γ-diversity. All diversity measures responded significantly positively to the ESBC treatment (Fig. 2B). However, effects were again stronger for taxonomic diversity than for phylogenetic diversity and stronger for q0 than q1 and q2 (Fig. 2B). The gain of species caused by ESBC treatments was on average ca. 40 (37-42) at a patch level (see Fig. S2-7).

Taxonomic β-diversity increased due to the ESBC treatment by 0.8% for q0 (Fig. 2C). This indicates that the positive effect of taxonomic γ-diversity for q0 is caused by both, an increase in α- and β-diversity. Taxonomic β-diversity for q1 showed a negative effect of ESBC of -1.4 %., however, the γ-diversity at q1 was still significantly higher. This indicates that the species gain for γ-diversity at q1 is mainly caused by the increase of α-diversity. Taxonomic β-diversity for q2 showed an increase of 1.4% and thus contributed to the positive effects of ESBC at γ-scale. Phylogenetic β-diversity for q0 was not significantly affected by the ESBC treatment but effects were significantly positive for q1 (0.7%) and negative for q2 (-1.8) (Fig. 2C, Fig. S2-S7).

Linear mixed effects models using α- and β-diversity metrics as response variables and deadwood volume and canopy openness as predictors revealed that canopy openness explains more of the diversity variation than deadwood volume in ESBC treatments (Table 1). Taxonomic α-diversity for q0 and q1 was significantly higher in open canopies in ESBC treatments. Phylogenetic α-diversity was significantly higher in open canopies in ESBC treatments only for q0. With exception of taxonomic β-diversity for q2, all β-diversity measures were significantly related to the canopy treatment in ESBC treatments (Table 1). For all measures of overall β-diversity (taxonomic, phylogenetic and all order of q), space was significant in control treatments. Low marginal R^2^s in comparison to the conditional R^2^s indicate high variability of effects among sites.

**Table 1.** Standardized effects sizes (t-value) from the models using fungal diversity measures (taxonomic, phylogenetic and taxonomic fungal functional lifestyle specific) as a response and deadwood volume and canopy openness as predictors. We used negative binomial models for taxonomic α-diversity and a gaussian linear mixed effects model for phylogenetic α-diversity and β-diversity (pairwise distances) considering site as a random effect (see Method section). We report the significance level (* p<0.05, ** p<0.01, *** p<0.001), the marginal (R^2^m) and the conditional (R^2^c) R-square.

| Model | Lifestyles |  |  | Main predictors |  |  |  | Covariates |  |  |  |  |  | Space |  | R<sup>2</sup>m/c |
| --- | --- | --- | --- | --- | --- | --- | --- | --- | --- | --- | --- | --- | --- | --- | --- | --- |
|  |  |  |  | Deadwood volume |  | Canopy openness |  | Ellenberg values Dim 1 (R~pH) |  | Ellenberg values Dim 2 (N) |  | Ellenberg values |  |  |  |  |
|  |  |  |  | C | ESBC | C | ESBC | C | ESBC | C | ESBC | C | ESBC | C | ESBC |  |
| $\alpha$ | Overall | TD | q0 | -0.23 | 1.69 | 1.91 | 3.54*** | 1.12 | 1.91 | 1.47 | 1.57 | - | | - | - | 0.10/0.70 |
|  |  |  | q1 | 0.66 | 1.40 | 1.48 | 2.87** | 1.29 | 1.84 | 0.32 | 0.08 | - | - | - | - | 0.08/0.59 |
|  |  |  | q2 | 1.06 | 1.57 | 1.50 | 1.27 | 1.45 | 1.52 | -0.42 | -0.89 | - | - | - | - | 0.07/0.32 |
|  |  | PD | q0 | -0.64 | 0.76 | 1.45 | 3.10** | 1.05 | 1.08 | 1.36 | 2.20* | - | - | - | - | 0.06/0.72 |
|  |  |  | q1 | 1.01 | 1.23 | 1.05 | 0.95 | 1.47 | -0.90 | -0.24 | 0.55 | - | - | - | - | 0.03/0.24 |
|  |  |  | q2 | 0.94 | 1.43 | 0.81 | -0.69 | 0.27 | -1.84 | -0.46 | 0.77 | - | - | - | - | 0.05/0.14 |
| $\beta$ | Overall | TD | q0 | 0.63 | 0.03 | 2.00* | 8.24*** | - | - | - | - | -0.40 | 2.84** | 4.88*** | 0.94 | 0.04/0.61 |
|  |  |  | q1 | 0.75 | 0.75 | -1.23 | 2.77** | - | - | - | - | -0.04 | 4.49*** | 6.43*** | -0.54 | 0.04/0.30 |
|  |  |  | q2 | 0.67 | 1.27 | -1.00 | 1.71 | - | - | - | - | -0.89 | 3.67*** | 4.60*** | -1.33 | 0.03/0.20 |
|  |  |  |  |  |  |  |  | - | - | - | - |  |  |  |  |  |
|  |  | PD | q0 | -0.06 | -1.51 | -1.61 | 4.96*** | - | - | - | - | -0.66 | 4.38*** | 4.26*** | -0.002 | 0.05/0.17 |
|  |  |  | q1 | -0.71 | 0.95 | -1.29 | 4.48*** | - | - | - | - | -0.04 | 2.46* | 3.94*** | -1.27 | 0.03/0.06 |
|  |  |  | q2 | -1.02 | 0.39 | -1.20 | 3.07** | - | - | - | - | -0.07 | 2.00* | 3.28** | -1.26 | 0.02/0.05 |
| $\alpha$ | Symbionts | TD | q0 | -0.44 | 2.11* | 0.04 | -1.66 | 1.52 | 2.69** | 0.31 | -0.05 | - | - | - | - | 0.08/0.46 |
|  |  |  | q1 | -0.20 | 0.31 | 0.63 | -0.31 | 1.77 | 3.04** | 0.70 | -1.11 | - | - | - | - | 0.11/0.31 |
|  |  |  | q2 | -0.20 | -0.71 | 0.50 | 0.37 | 1.32 | 2.31 | 0.67 | -1.43 | - | - | - | - | 0.08/0.21 |
|  |  |  |  |  |  |  |  |  |  |  |  | - | - | - | - |  |
|  | Saprotrophs |  | q0 | 0.15 | 1.98* | 2.31* | 2.79** | 1.36 | 2.25* | 0.74 | 1.35 | - | - | - | - | 0.12/0.55 |
|  |  |  | q1 | 0.35 | 2.81** | 1.18 | 0.29 | 1.60 | 1.64 | -1.58 | 0.67 | - | - | - | - | 0.08/0.54 |
|  |  |  | q2 | 0.13 | 2.96** | 0.66 | -1.95 | 1.71 | 1.42 | -2.14* | -0.04 | - | - | - | - | 0.10/0.35 |
|  | Parasites |  | q0 | -0.37 | 1.06 | 2.64** | 3.06** | 2.56* | 3.99*** | 0.26 | -0.56 | - | - | - | - | 0.29/0.42 |
|  |  |  | q1 | 0.85 | 0.60 | 0.25 | 1.62 | 2.66** | 3.40*** | 0.49 | -0.74 | - | - | - | - | 0.20/0.46 |
|  |  |  | q2 | 0.54 | 0.25 | 0.12 | 1.38 | 2.53* | 3.38*** | -0.36 | -0.90 | - | - | - | - | 0.20/0.36 |
| $\beta$ | Symbionts | TD | q0 | 0.85 | 0.90 | 2.45* | 5.69*** | - | - | - | - | 0.19 | 3.38*** | 4.97*** | 0.17 | 0.02/0.62 |
|  |  |  | q1 | 0.17 | 1.39 | 0.26 | 6.05*** | - | - | - | - | -1.72 | 3.78*** | 4.12*** | -0.56 | 0.05/0.43 |
|  |  |  | q2 | -0.26 | 1.54 | -0.61 | 4.54*** | - | - | - | - | -2.25* | 1.90 | 3.33*** | -0.68 | 0.04/0.26 |
|  | Saprotrophs |  | q0 | -0.92 | -0.23 | 0.79 | 4.23*** | - | - | - | - | -0.77 | 2.53* | 2.81** | 1.49 | 0.02/0.54 |
|  |  |  | q1 | -0.19 | 0.80 | 0.38 | 3.69*** | - | - | - | - | -0.40 | 0.35 | 4.95*** | 1.67 | 0.02/0.30 |
|  |  |  | q2 | -0.11 | 1.58 | 0.58 | 2.08* | - | - | - | - | -0.92 | -1.05 | 3.59*** | 0.29 | 0.02/0.16 |
|  | Parasites |  | q0 | -1.01 | 0.29 | -0.05 | 2.24* | - | - | - | - | 2.90* | 2.09* | 2.86** | -0.48 | 0.02/0.23 |
|  |  |  | q1 | 1.06 | -1.25 | -2.76** | 6.47*** | - | - | - | - | 0.84 | 3.66*** | 4.85*** | -1.47 | 0.06/0.19 |
|  |  |  | q2 | 0.98 | -1.60 | -3.09** | 4.10*** | - | - | - | - | -0.18 | 2.88** | 3.10** | -2.78** | 0.04/0.13 |

### Functional lifestyles

The α-, β- and γ-diversity of symbiotic fungi decreased significantly with the ESBC treatment for q0 (Fig. 2 D-F, Fig. S8-10). γ-diversity decreased by 8 (7.6-8.7) species. Further, the ESBC treatments decreased the symbiont γ- and α-diversity for q1 and q2 (but not significantly for q2 for γ-diversity). In contrast symbiont β-diversity for q1 and q2 increased by 3.4% and 4.8% in ESBC treatments. However, positive response of symbiont β-diversity did not result in significant positive γ-effects.

For saprotrophic fungi, α-, β- and γ-diversity increased significantly with the ESBC treatments for all orders of q (Fig. 2 D, E, Fig. S11-13). Effects were strongest for q0 causing a gain of species of ca. 54 (47-62) and 25 (24-27) at γ- and α-scale respectively. β-diversity for all q orders was significantly lower in ESBC treatments with strongest effects on q2 (-4.8%). However, negative β-effects did not out cancel overall positive effects at γ-scale.

For parasitic fungi, γ-diversity at q0 was not significantly related to the ESBC treatments (Fig. 2D, Fig. S14). Parasitic α-diversity at q0 was positively (gain of species by 7; 5.7-8.7) and β-diversity negatively related to the ESBC treatments (-2%,) (Fig. 2E, F). This contrast between α- and β-diversity might cause a non-significant γ-effect. For q1 and q2, parasitic α- and γ-diversity was negatively related to ESBC treatments, however, effects were not strong (Fig. 2D, E, Fig. S15, 16). For parasitic β-diversity, effects of the ESBC treatments were positive for q1 (1.8%) and q2 (4.8%) (Fig. 2C).

Linear mixed effects models using α- and β-diversity metrics as response and deadwood volume and canopy openness as predictors separately for the functional lifestyles revealed contrasting effects (Table 1). Symbiont diversity showed weak responses to the predictors in both control and ESBC treatments. While we observed a significant positive relationship between symbiont α-diversity for q0 with deadwood amount in the ESBC treatments, effect size of canopy openness was negative (but not significant, Table 1). Saprotrophic α-diversity for all q orders was significantly positive related to deadwood amount in the ESBC treatments (Table 1). Further, we found a significant positive relationship between saprotrophic α-diversity for q0 and canopy openness in control and ESBC treatments. α-diversity of parasites for q0 were significantly positive related to canopy openness in control and ESBC treatments. β-diversity of all functional lifestyles were significantly related to canopy openness across orders of q (Table 1). Across all order of q and lifestyles, we revealed a significant effect of space on β-diversity in control treatments (Table 1).

## Discussion

By comparing homogenized versus experimentally structurally enhanced forests (ESBC) replicated across Germany, we revealed that ESBC increased fungal γ-diversity at landscape scale as expected. However, in contrast to our expectation, this effect was mainly caused via species gains at α-forest stand scale. Further, species gain is mainly caused by canopy openness and not resource availability as hypothesized. Weak response of phylogenetic diversity to ESBC indicates that the species gain comes from lineages already existing in the landscape supporting the *functional similarity hypothesis*. Further, we revealed contrasting effects and determinants of ESBC on the fungal functional lifestyles across scales.

According to our expectation, we found that ESBC increased fungal taxonomic diversity at the γ-scale for all orders of q. The increase of species diversity was consistent at α- and β- level for q0 and q2. For β-diversity of q1, we revealed a negative effect of ESBC, but this did not cause a negative effect on γ-diversity. From these findings, we overall conclude that the species gain at landscape scale is mainly caused by the positive effects observed at α-scale. This contrasts with our expectation, as we expected a dominating effect of ESBC on β-diversity rather than α-diversity. Numerous studies demonstrated consistent effects of environmental variables on soil fungal communities at different spatial scales (e.g., Tedersoo et al. 2014, Mikryukov et al. 2023, Odriozola et al. 2024). However, what we learn from our study is that the strong effects of β-diversity observed in previous studies at local scale, might not necessarily translate to increase larger scale γ-diversity. Nevertheless, beside the strong effect of α-diversity observed in our study, we also revealed an independent contribution of β-diversity explaining the increased γ-diversity in ESBC forests.

We expected that the environmental factors manipulated in the ESBC treatment would affect α- and β-diversity components in different ways. We hypothesized that an increase in α-diversity is related to an increase in resource availability caused by the deadwood enrichment which would be in line with the *more individual hypothesis* (MacArthur and MacArthur 1961). However, in the overall models, canopy openness turned out to be more important. We expected that the deadwood increase should affect soil chemical and physical properties via decomposition processes and changes in nutrition sources. For example, in one study it has been shown that availability of deadwood increases soil organic carbon, total nitrogen content and exchangeable cations (Sokołowski et al. 2025). In 8 out of the 11 sites, deadwood has been created in 2016 respectively 2018 and thus, deadwood decomposition should be pronounced (Schreiber et al. 2024) with effects on soil properties. One explanation for the non-significant relationship between overall diversity and deadwood enrichment might be that the effects of the decomposition with subsequent nutrient release is locally restricted and patchy and thus, did not reach a critical amount across the spatial scale of the forest stand. Hence, the small amount of soil sampled for fungal metabarcoding might be not sufficiently enriched with nutrients from the deadwood decay at patch scale. However, we found significant positive effects of deadwood enrichment on saprotrophic diversity indicating that contrasting effects among functional lifestyles might out cancel overall effects (see discussion below). Nevertheless, mores studies are needed measuring the amount and spatial structure of carbon and nutrient input by deadwood over the course of succession across the patch scale.

Overall α-diversity was positively related to canopy openness in the ESBC treatments. It has been shown that open forest canopies increase microclimate extremes but also increase the average temperature (Thom et al. 2020, Schreiber et al. 2025b). From our findings, we conclude that microclimate harshness does not act as a filter causing a reduction in fungal diversity. Instead, the increase in mean annual temperature in open canopies might increase metabolic rates, thereby allowing more individuals and thus species to coexist (Hawkins et al. 2007, Schreiber et al. 2025a). Another explanation might be that the microclimate niches increase in a patch under open canopies compared to closed canopies (Thom et al. 2020). Thus, increased diversity under open canopies might be explained by the habitat heterogeneity hypothesis. This would be in line with a larger-scale study demonstrating that fungal species richness increase in areas with a high seasonal variability in temperature conditions (continentality) (Bässler et al. 2022). We found further that canopy openness with ESBC treatments affected significantly the community composition as expected which is in line with several studies (e.g., Castaño et al. 2018, Rodriguez-Ramos et al. 2021). However, as outlined above, we found that the contribution of β-diversity on γ-diversity is less pronounced than effects of α-diversity.

We found significant effects of space affecting β-diversity in control but not in ESBC treatments. This suggests that the metacommunity dynamics and thus, assembly mechanisms might differ between homogenous and heterogenous forests. From theory, homogeneous environmental conditions support metacommunities dominated by patch dynamics (Leibold et al. 2004). Thus, we suggest that deterministic assembly processes like species sorting are more prevalent in ESBC than in control treatments. In the homogenous control forests, other processes like dispersal or drift might prevail. We do not assume that dispersal limitation is relevant for fungi at the scale of our study (within the landscapes) (Abrego et al. 2018, Komonen and Müller 2018). However, homogenizing dispersal (i.e., mass effects) or drift might explain our observed effect of space and thus, contribute to explain the dominant assembly process in homogenous forests. Further studies are needed applying rigorous null models to test whether and how ESBC change species assembly processes in comparison to homogenous forest ecosystems.

Taxonomic diversity showed stronger responses to ESBC treatments than phylogenetic diversity. Further, effects were stronger for q0 than q1 and q2. This indicates that the gain of species caused by ESBC at landscape scale is particularly related to rare species from lineages already existing in the landscape. Phylogenetic diversity has been suggested as a measure of functional diversity if functional traits are conserved within major lineages (Srivastava et al. 2012), but this assumption has been theoretically and empirically challenged (Winter et al. 2013, Mazel et al. 2018). Note that measuring functional traits for 20.000 putative fungal species (OTUs) is currently impossible and more generally, our understanding about functionally relevant traits at species level is limited given the tremendously high diversity of species (Dawson et al. 2018). At least, we can conclude that a change in environmental conditions by forest management strategies as reflected by our ESBC treatments does not have substantial effects on the major fungal lineages at landscape scale. We, therefore, cautiously suggest that the gain of species by ESBC might not necessarily add further functions to the system. Overall, this would support the *functional similarity hypothesis* and not the *insurance hypothesis*.

By analyzing the response of the diversity of the main fungal functional lifestyles separately in relation to the ESBC treatment, we revealed contrasting patterns. This contrast becomes most obvious when focusing on the response of the diversity of q0 at γ-scale. While symbionts showed a significant negative effect with ESBC, saprotrophic diversity increased considerably, and parasitic fungi showed no effect. Our environmentally informed models partly help to explain these findings. First, the diversity of symbiont fungi revealed mostly non-significant relationships with deadwood volume and canopy openness. One exception is the significant increase of symbiont richness (q0) with deadwood volume. However, it seems that our predictors cannot sufficiently explain the overall negative response of the symbiont diversity. One variable missing that might explain the negative relationship, is the reduced amount of host tree species with our ESBC manipulation (ca. 30% of trees removed). Thus, logging activities of the main host trees could reduce niche space and, therefore, symbiotic diversity, which has been also shown in other studies (Tomao et al. 2020). Note that most of the symbiont species sampled in our setting were ectomycorrhizal fungi. As our forest system is mainly characterized by obligate ectomycorrhizal associations (e.g., oak, beech Müller et al. 2023), we suggest that the observed negative relationship is mainly caused by the reduced amount of host resources (e.g., root space). Further studies are needed, whether and if so at which temporal scale the observed effect might diminish caused by increased growth response of the remaining trees (with associated root increase) and/or due to the recolonization of young ectomycorrhizal seedlings. Second, the gain in saprotrophic diversity by ESBC seemed to be driven by both deadwood volume and canopy openness, both showing significant positive effects. Mechanisms explaining the diversity increase of canopy openness might be the same as discussed above for overall diversity response. However, the consistent significant positive response of deadwood volume on saprotrophic diversity supports the assumption that deadwood enrichment increases resource availability for soil fungi. Thus, for saprotrophic fungi, we found support for the *more individuals hypothesis*. Third, parasitic diversity did not respond to ESBC at landscape γ-scale but showed significant positive effects of canopy openness for q0 α-diversity. As also discussed for saprotrophic diversity, metabolic opportunities could also be important here. Whether an increase of pathogen richness is also caused by an increase of host species (e.g., herb layer diversity) or whether an increase in pathogen richness increase pathogen severity for their hosts with consequences on diversity dynamics must be left to further studies.

## Conclusions

Our spatially explicit and replicated experiment on the effect of heterogeneity in temperate forests provides robust empirical evidence that any enhancement of between patch heterogeneity by light and deadwood increases the landscape wide diversity of soil fungi. However, as the response vary across scales and functional groups, our results provide further insights for the restoration of forests structure. At the local scale saprotrophic fungi diversity increased with increasing amount of deadwood and in canopy gaps, while symbiotic fungi diversity of rare species tends to decrease in the gaps. In contrast to these various α-effects, β-diversity consistently increased with increasing heterogeneity in canopy cover for all functional groups and Hill numbers. This suggests that aiming on restoration of the various functional groups of this cryptic kingdom of fungi in forests soils require a highly heterogenous mosaic of dense and open forests and variation in deadwood amount, while the current homogenous, mature production forests promote mainly symbiotic fungi closely related to the growing stock of trees.

## Acknowledgements

We thank Michael Junginger, and all assistants in the field and laboratory. We also thank the entire BETA-FOR team for their support. This work was financed by the Deutsche Forschungsgemeinschaft [BETA-FOR, 459717468].

